# Cortical idiosyncrasies predict the perception of object size

**DOI:** 10.1101/026989

**Authors:** Christina Moutsiana, Benjamin de Haas, Andriani Papageorgiou, Jelle A. van Dijk, Annika Balraj, John A. Greenwood, D. Samuel Schwarzkopf

## Abstract

Perception is subjective. Even basic judgments, like those of visual object size, vary substantially between observers and also across the visual field within the same observer. The way in which the visual system determines the size of objects remains unclear, however. We hypothesize that object size is inferred from neuronal population activity in V1 and predict that idiosyncrasies in cortical functional architecture should therefore explain individual differences in size judgments. Indeed, using novel behavioral methods and functional magnetic resonance imaging (fMRI) we demonstrate that biases in size perception are correlated with the spatial tuning of neuronal populations in healthy volunteers. To explain this relationship, we formulate a population read-out model that directly links the spatial distribution of V1 representations to our perceptual experience of visual size. Altogether, we suggest that the individual perception of simple stimuli is warped by idiosyncrasies in visual cortical organization.

---

How do we perceive the size of an object? A range of recent observations have lent support to the hypothesis that the visual system generates the perceived size of an object from its cortical representation in early visual cortex^1^. In particular, the spatial spread of neural activity in visual cortex has been related to apparent size under a range of contextual modulations^2–7^. The strength of contextual size illusions has further been linked to the cortical territory in V1 that represents the central visual field^8,9^. These findings suggest that lateral connections in V1 may play a central role in size judgments because these interactions are reduced when V1 surface area is larger. Indeed, similar interactions have been argued to underlie the strength of the tilt illusion^10,11^, perceptual alternations in binocular rivalry^12^, the influence of distractors in visual search tasks^13^, and visual working memory capacity^14^. Even the precision of mental imagery co-varies with V1 area^15^ suggesting V1 may be used as a ‘workspace’ for storing mental images whose resolution is better when surface area is larger.

However, these previous findings do not demonstrate that V1 representations *per se* are relevant for size judgments, and in particular for *subjective* judgments of object size. If V1 signals were indeed the basis for these judgments then variations in the functional architecture of V1 should explain *idiosyncratic biases in basic size perception* (i.e. size judgements that occur in the absence of any contextual/illusory effects). To date this prediction remains untested. Previous neuroimaging experiments have focused on modulations of apparent size that must involve additional processing, either due to local interactions between adjacent stimuli in V1 or by a context that likely involves processing in higher visual areas. Others have shown that the *objective* ability to discriminate subtle differences between stimuli is related to cortical magnification and spatial tuning in early visual cortex^11,16,17^. However, no experiment to date has shown a relationship between V1 and *subjective* perceptual biases in the *absence of any contextual interaction*, even though there are considerable individual differences in perceptual biases.

It is well established that subjective size judgments for simple, small stimuli can vary substantially between observers and even across the visual field within the same observer. Previous behavioral research has shown that small visual stimuli appear smaller when they are presented in the periphery^18–20^. A simple explanation for this could be the impoverished encoding of stimuli in peripheral vision. However, when stimuli are dimmed artificially to mimic the peripheral decrease in visibility, the same biases are not found^19^. Another explanation could be that higher brain regions that integrate the perceptual input to V1 into a behavioral decision are poorly calibrated against the decrease in cortical magnification when moving from central to peripheral vision. Small errors in this calibration would cause a residual misestimation of stimulus size based on V1 representations and in turn lead to perceptual misestimation^20^. However, neither of these models can explain why these perceptual biases are consistent underestimates of stimulus size. Impoverished stimulus encoding alone should only result in poorer acuity while residual errors in calibration would be expected to show both under- and overestimation. Furthermore, recent research has also demonstrated reliable heterogeneity in size judgments across the visual field within individual observers at iso-eccentric locations^9,21^. We can consider these variations as a ‘perceptual fingerprint’ that is unique to each observer. The neural basis of these individual differences however remains unknown.

In the present study we used fMRI to compare perceptual biases in size judgments with individual functional architecture in V1 – specifically, the population receptive field (pRF) spread and local cortical surface area. To do so we developed the Multiple Alternative Perceptual Search (MAPS) task. This approach combines a matching task with analyses similar to reverse correlation^22,23^. Observers search a peripheral array of multiple candidate stimuli for the one whose subjective appearance matches that of a centrally presented reference. This task allows measurement of subjective appearance whilst minimizing the decisional confounds present in more traditional tasks like stimulus adjustment or the method of constant stimuli^24–26^. The MAPS task further estimates perceptual biases and discrimination acuity while several stimuli are presented simultaneously. We consider this a more naturalistic task, akin to our daily perceptual judgments (Figure 1A), compared with traditional psychophysical tasks involving single, isolated objects.

**Figure 1.**
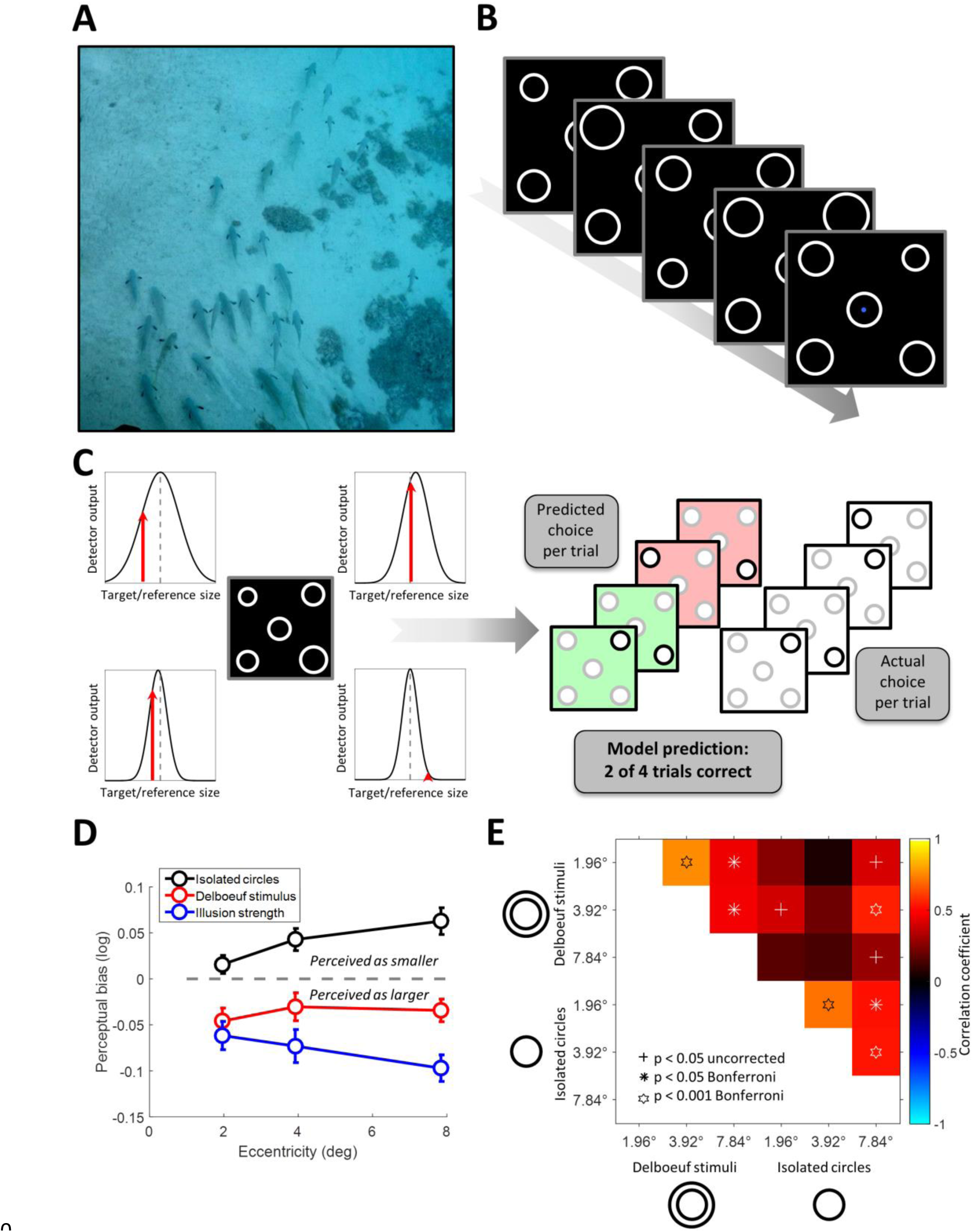
Idiosyncratic biases in size perception. **A**. Visual objects often appear in the presence of similar objects. For example, a spearfisherman may be searching this school for the largest fish. What is the neural basis for this judgment? **B**. The MAPS task. In each trial, observers fixated on the center of the screen and viewed an array of five circles for 200ms. The central circle was constant in size, while the others varied across trials. Each frame here represents the stimulus from one trial. The arrow denotes the flow of time. Observers judged which of the circles in the four corners appeared most similar in size to the central one. **C**. Analysis of behavioral data from MAPS task. The behavioral responses in each trial were modeled by an array of four “neural detectors” tuned to stimulus size (expressed as the binary logarithm of the ratio between the target and the reference circle diameters). Tuning was modeled as a Gaussian curve. The detector showing the strongest output to the stimulus (indicated by the red arrows) determined the predicted behavioral response in each trial (here, the top-right detector would win). Model fitting minimized the prediction error (in this example the model predicted the actual behavioral choice correctly for 50% of trials) across the experimental run by adapting the mean and dispersion of each detector. **D**. Average perceptual bias (positive and negative: target appears smaller or larger than reference, respectively), across individuals plotted against target eccentricity for simple isolated circles (black), contextual Delboeuf stimuli (red), and relative illusion strength (blue), that is, the difference in biases measured for the two stimulus conditions. Error bars denote ±1 standard error of the mean. **E**. Correlation matrix showing the relationship of unique patterns of perceptual biases in the two conditions (isolated circles and Delboeuf stimuli) and at the three target eccentricities. Color code denotes the correlation coefficient. Symbols denote statistical significance. Crosses: p<0.05 uncorrected. Asterisks: p<0.05 Bonferroni corrected. Hexagrams: p<0.001 Bonferroni corrected.

## Results

Thirteen normal, healthy observers viewed an array of 5 circles on each trial and made a perceptual judgment (Figure 1B). The central circle was constant in size and served as the reference. Observers reported which of the four target circles appeared most similar in size to the reference. We fit a model to explain each observer’s behavioral responses, with each of the four target locations modeled via the output of a detector tuned to stimulus size. In each trial the detector showing the strongest response was used to predict the observer’s behavioral choice. This procedure allowed the estimation of both raw perceptual bias and uncertainty (dispersion) at each location (Figure 1C).

### Apparent size depends on eccentricity

Peripheral stimuli appeared smaller on average than the central reference, confirming earlier reports^18,20,27^. This reduction in apparent size increased with stimulus eccentricity (Figure 1D, black curve). When instead of isolated circles we presented the target circles inside larger concentric circles, perceptual biases were predictably shifted in the other direction (the Delboeuf illusion^28^) so that targets appeared on average larger than the reference (Figure 1D, red curve). This illusory effect interacted with the effect of eccentricity on apparent size, leading again to a gradual reduction in (illusory) size as stimuli moved into the periphery. This differs somewhat from the classical Delboeuf illusion, where perceptual biases are typically compared to a reference either at the same eccentricity or even at the same stimulus location. In contrast, in our task the reference is at fixation. To disentangle the illusion from the effect of eccentricity, we therefore also calculated the *Relative illusion strength*, that is, the difference in perceptual bias for isolated circles and the illusion stimuli at each location. This effect (here an *increase* in apparent size) also increased with eccentricity (Figure 1D blue curve; but note that since observers never compared the stimuli directly at iso-eccentric locations this may not fully account for the classical Delboeuf illusion). To summarize, objects appear increasingly smaller as they move into peripheral vision, where the magnitude of size illusions also has an increasing effect (here with the Delboeuf illusion to make them appear larger).

These results cannot be trivially explained by differences in discrimination acuity. Because spatial resolution decreases in peripheral vision, it is theoretically possible that bias estimates are noisier at greater eccentricities and thus produce this pattern of results, in particular for the Delboeuf stimuli where bias magnitude decreases. To rule out this confound we also calculated mean bias estimates weighted by the precision of observers’ size estimates (i.e. the reciprocal of dispersion) at the corresponding locations. The pattern of results is very similar to the one for raw biases (Supplementary Figure S1A). Two years after the initial experiment, we also conducted another small experiment on four observers. In this experiment we included two larger eccentricities (11.76° and 15.68°). This confirmed that the size of isolated circles continue to be underestimated even at larger eccentricities. In contrast, although the size of Delboeuf stimuli is overestimated at the more central eccentricities, the bias magnitude decreases with eccentricity and is close to zero at the most peripheral location tested (Supplementary Figure S1B). However, the accuracy of performing the MAPS task also decreases with eccentricity, especially for Delboeuf stimuli (Supplementary Figure S1C) presumably because crowding makes it difficult to separate the inner and outer circle.

### Idiosyncratic biases in size perception

Critically, we next analyzed the *idiosyncratic pattern* of perceptual biases for each observer by comparing biases across the visual field, for both isolated circles and Delboeuf stimuli. To do so, bias estimates were taken from all observers in each visual field quadrant, separately for each eccentricity and stimulus type (that is, each visual field location was treated as a separate data point, so n=40). Biases were strongly correlated across both stimulus type and over the three eccentricities tested (Figure 1E and Supplementary Figure S2). That is, if observers perceived a strong reduction in the apparent size of a stimulus at a given location, they tended to show strong reductions for the same stimulus type within the same visual field quadrant, regardless of eccentricity. Variations between different quadrants of this kind are consistent with the anatomical separation of visual quadrant maps in retinotopic areas. Psychophysical studies suggest that similarly coarse differences may be common, for instance with the frequent observation that performance in the lower visual field exceeds that in the upper visual field^29,30^. We further confirmed that these bias estimates were highly reliable even between testing sessions separated by one or two years (Supplementary Information: Reliability of perceptual bias estimates).

### Perceptual biases correlate with spatial tuning in visual cortex

Next we employed functional magnetic resonance imaging (fMRI) with population receptive field (pRF) mapping to estimate the tuning of V1 voxels to spatial position (Figure 2A–B^31–33^). Importantly, these neuroimaging experiments were independent from the behavioral experiments and conducted many months later in a different testing environment (MRI scanner vs behavioral testing room).

**Figure 2.**
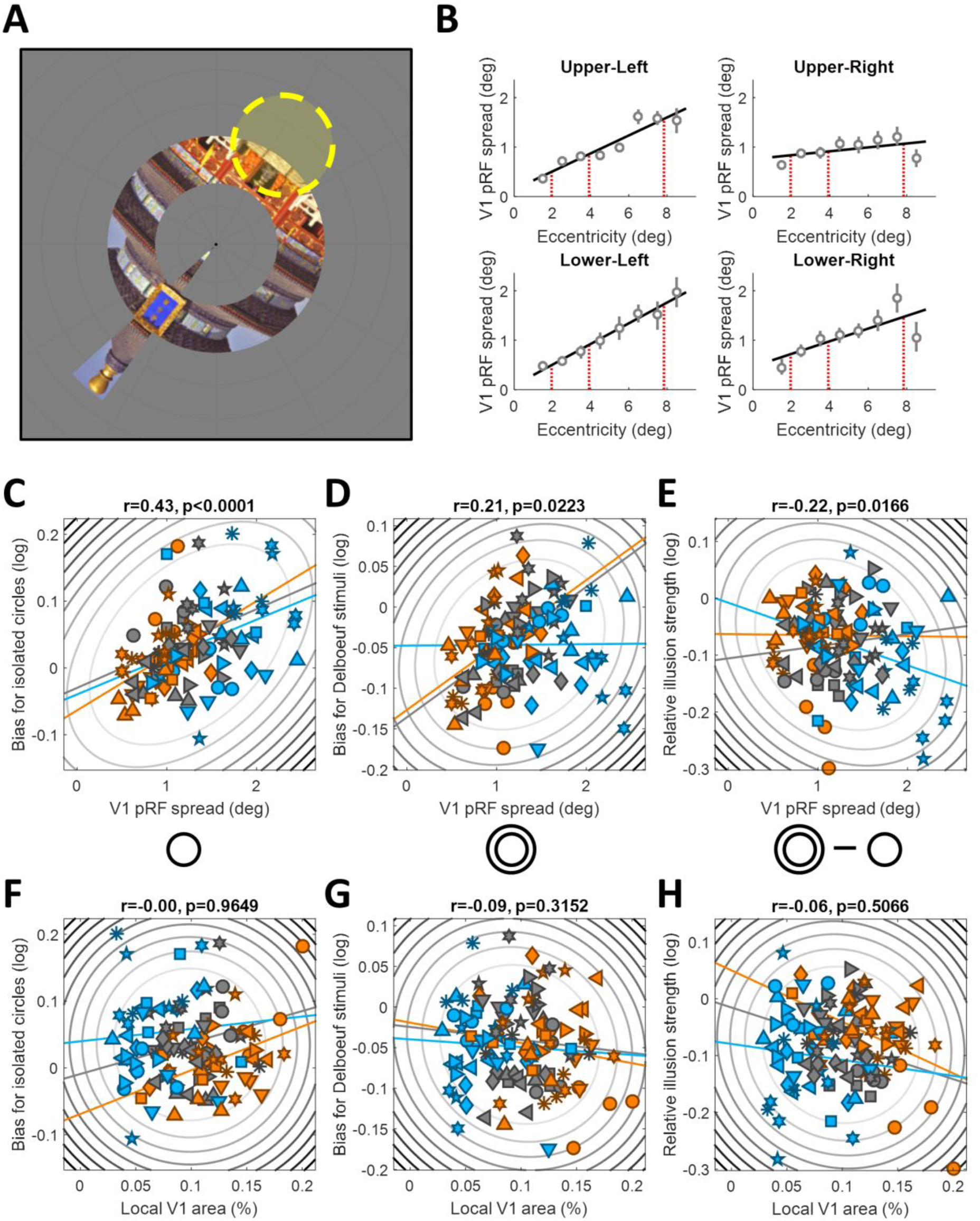
Neural correlates of size perception. **A**. Population receptive field (pRF) mapping with fMRI. Observers viewed natural images presented every 500ms within a combined wedge-and-ring aperture. In alternating runs the wedge rotated clockwise/counterclockwise in discrete steps (1Hz) around the fixation dot while the ring either expanded or contracted. A forward model estimated the position and size of the pRF (indicated by yellow circle) that best explained the fMRI response to the mapping stimulus. **B**. We estimated the pRF spread corresponding to each target location in the behavioral experiment by fitting a first-order polynomial function (solid black line) to pRF spreads averaged within each eccentricity band for each visual quadrant and extrapolating the pRF spread at the target eccentricities. Grey symbols in the four plots show the pRF spread by eccentricity plots for the four target locations (see insets) in one observer. Grey error bars denote bootstrapped 95% confidence intervals. The solid black line shows the polynomial fit used to estimate pRF spread at each target location. The vertical red dashed lines denote the three stimulus eccentricities at which we extrapolated pRF spread and surface area from the fitted polynomial functions. Data from other observers and V1 surface area measurements are included as Supplementary Information. **C-E**. Perceptual biases for isolated circles (**C**), Delboeuf stimuli (**D**), and the relative illusion strength (**E**) plotted against pRF spread for each stimulus location and observer. **F-H**. Perceptual biases for isolated circles (**F**), Delboeuf stimuli (**G**), and the relative illusion strength (**H**) plotted against V1 surface area (as percentage of the area of the whole cortical hemisphere) for each stimulus location and observer. In **C-I** Symbols denote individual observers. Elliptic contours denote the Mahalanobis distance from the bivariate mean. The colored, straight lines denote the linear regression separately for each eccentricity. Colors denote stimuli at 1.96° (orange), 3.92° (grey), or 7.84° (light blue) eccentricity.

Interestingly, this analysis revealed a systematic relationship between perceptual biases and pRF spread (also known as pRF size or the σ parameter of the Gaussian pRF model). With data averaged across observers, increasing eccentricity gives both an increase in pRF spread^31^ (see Supplementary Data File 1) and a decrease in apparent size (Figure 1D). We then considered individual data by calculating the correlation between pRF spread and perceptual biases. To do so, we considered every stimulus *location* in every observer as a separate observation (n=120). Both isolated circles (r=0.43, p<0.0001; Figure 2C) and Delboeuf stimuli (r=0.21, p=0.0223; Figure 2D) were perceived as smaller when they were presented at visual field locations covered by voxels with larger pRFs. We obtained similar results when analyzing data separately for each eccentricity (n=40), except for the Delboeuf stimuli at the largest eccentricity (Supplementary Figure S3A–B). These individual differences demonstrate that there is correlation between pRFs and apparent size for *idiosyncratic variations* at a *fixed* eccentricity. Our relative illusion strength also showed a negative correlation with pRF spread (r=-0.22, p=0.0166, n=120; Figure 2E), indicating that larger pRFs were associated with smaller differences between raw biases for Delboeuf stimuli and isolated circles. This result was however largely driven by the results for the largest eccentricity (Supplementary Figure S3C).

The most critical of the above tests of our hypothesis that cortical properties and perception are linked treated each visual field location as a separate observation. While this includes the between-subject variance as well as the pattern of differences within each observer, it directly quantifies the relationship between the two variables. However, the measurements from the four visual field quadrants for a given observer are naturally not independent. We therefore conducted several additional tests. We first repeated all of these analyses after subtracting the mean bias/pRF spread from each observer and eccentricity. This allows analysis of the pattern of results across quadrants whilst removing both the individual differences between observers (between-subject variance) and differences related to eccentricity. This analysis confirmed the correlation (n=120) between pRF spread and biases for isolated circles (r=0.29, p=0.001). For Delboeuf stimuli and the relative illusion strength the correlations were not significant, though they showed the same trends as the equivalent correlations in the main analysis (Delboeuf stimuli: r=0.15, p=0.112; illusion strength: r=-0.16, p=0.077). We also conducted a similar second-level analysis in which we first calculated the correlations across the four locations separately in each observer at each eccentricity and then tested whether the average correlation (after z-transformation) was significantly different from zero. Finally, we performed a multivariate canonical correlation analysis using the four observations per observer and eccentricity (see Supplementary Information: Intra-individual differences analysis for more detail on the different analyses). The results of these additional analyses are shown in Table 1.

**Table 1.**
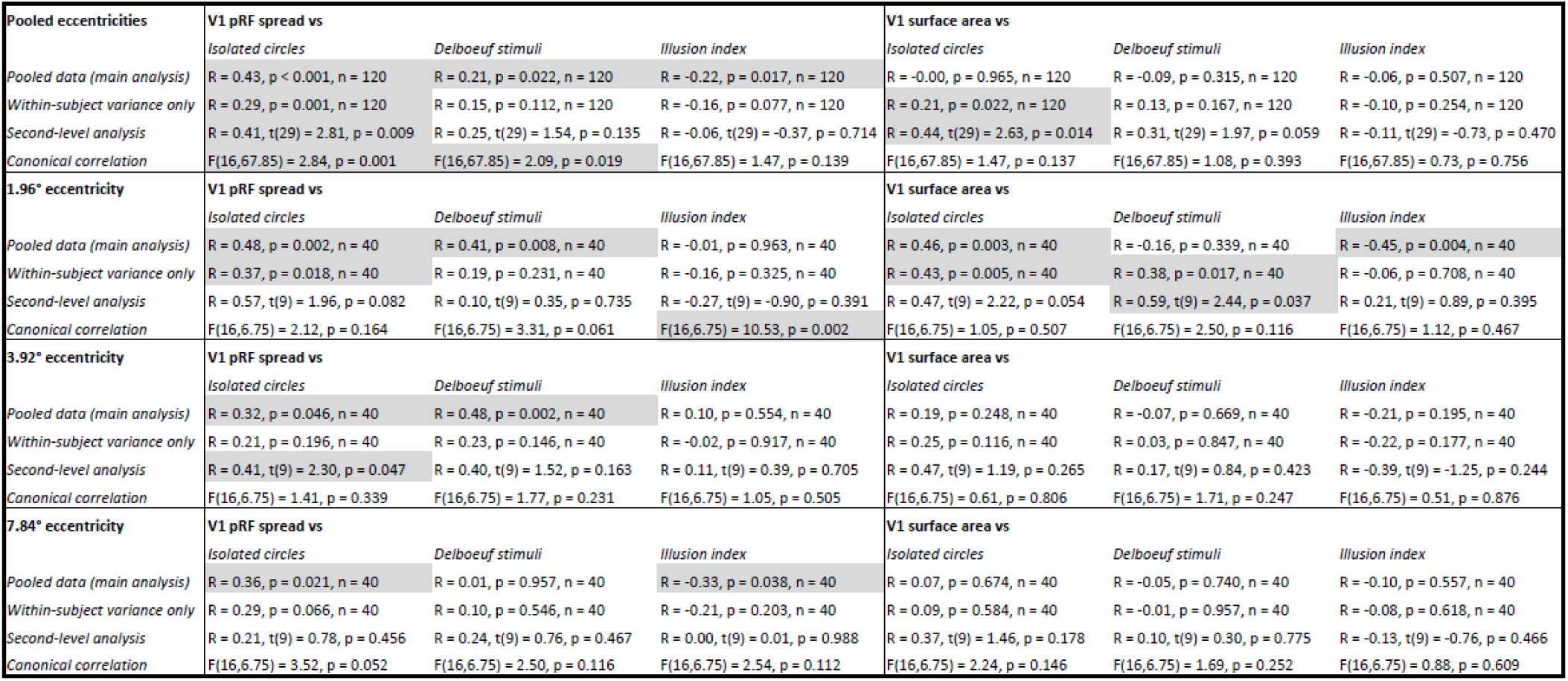
All correlations between pRF spread or cortical surface area in V1 and perceptual bias measures using four complementary analysis approaches: *Pooled data* refers to the main analysis presented in which we simply treated each of the 12 visual field locations per observer as an separate data point. *Within-subject variance only* refers to the equivalent analysis after removing the mean pRF spread, surface area and perceptual bias across the four locations for each eccentricity and observer. Second-level analysis refers to the analysis in which we first calculated the correlation across four locations separately for each observer and eccentricity and then determined whether the average correlation (after z-transformation) was different from zero. Both, the average correlation coefficient and the statistics of the t-test against zero are shown. *Canonical correlation* corresponds to a standard canonical correlation analysis. This is expressed as an F-test. Only the full combination of all four visual field locations per observer and eccentricity are used. Across the table, cells shaded in grey denote correlations statistically significant at p<0.05.

A similar approach exploiting within-subject correlations has previously been used in the context of retinotopic mapping data^34,35^ and spatial heterogeneity in perceptual function^21^. These studies suggests that our sample size of 10 observers is likely sufficient. However, to confirm this we also performed a simulation analysis to quantify the statistical power of our approach. Our main analysis and the one removing between-subject variance had the greatest sensitivity for detecting a true effect (with approximate power of 90% for an assumed true correlation of r=0.3). This is unsurprising given the large number of data points in these analyses (n=120). However, while all other analyses produced nominal false positive rates of ∼5%, false positives rose slightly to ∼9% when removing between-subject variance. This suggests our main analysis as the optimal statistical test for our hypothesis, affording high sensitivity and specificity (see Supplementary Information: Power analysis).

Finally, we also analyzed the equivalent correlations between behavioral measures and pRF spread for areas V2 and V3. Pooled across eccentricities (n=120) pRF spreads in either area were significantly correlated with the biases for isolated circles (V2: r=0.4, p<0.0001; V3: r=0.29, p=0.0013). Correlations between pRF spread in these areas and the biases for Delboeuf stimuli followed the same trend but were not significant (V2: r=0.14, p=0.1188; V3: r=0.16, p=0.0728). Moreover, separated by eccentricity all of these correlations were positive but not significant. Thus the relationship between pRF spread and perceptual biases was not specific to V1 but a general feature of early visual cortex. This is unsurprising given that pRF spreads in V1 were largely well correlated with the extrastriate regions (minimal correlation, separately for each eccentricity (n=40) in V2: r=0.49, p=0.0014; V3: r=0.26, p=0.1037).

### Basic read-out model of size perception

Why should the apparent size of our circle stimuli be smaller when pRFs are larger? While this result is consistent with the simple impoverishment hypothesis, which states that perceptual biases depend on the precision of the stimulus representation, this alone does not explain why biases are consistent *underestimates* of stimulus size^19^. To understand this better we conducted a series of simulations that assume that higher brain regions involved in integrating sensory inputs into a perceptual decision about object size read out signals from V1 neurons^36^. We simulated the neuronal activity inside the retinotopic map by passing the actual spatial position of the two edges of the stimulus through a Gaussian filter bank covering that stimulus location. The stimuli were simulated as a binary vector representing 1050 pixels (corresponding to the height of the screen) where the edges were set to 1 while the background was set to 0. The filter bank assumed a Gaussian tuning curve whose width was parameterized at each pixel along this vector. We calculated the response of each filter, to give rise to a population activity profile. Subsequently, a higher level then sampled this activity to infer stimulus size. This basic model only assumes two layers – however, it is principally the same if activity is submitted from V1 across multiple stages along the visual hierarchy with each layer applying similar filtering.

With this approach, stimulus size can be inferred from the distance between the activity peaks corresponding to the two edges (Figure 3). When the spatial tuning of visual neurons (i.e. neuronal receptive field size) is narrow, the peaks can accurately localize the actual stimulus edges. However, as tuning width increases, the activity profile becomes blurrier. Critically, the distance between the two peaks also becomes *smaller* because activity from the two edges is conflated (Figure 3A). It follows that with wider tuning (at greater eccentricity and larger pRF spread), estimates of the separation between peaks decreases until eventually the two peaks merge. This scenario presumably corresponds to far peripheral vision. For the Delboeuf stimuli, the separation of peaks is greater than that of the actual stimulus edges when tuning width is narrow because activity corresponding to the inducer and the target blurs together. However, as tuning width increases the separation also becomes smaller just as for isolated circles (Figure 3B).

**Figure 3.**
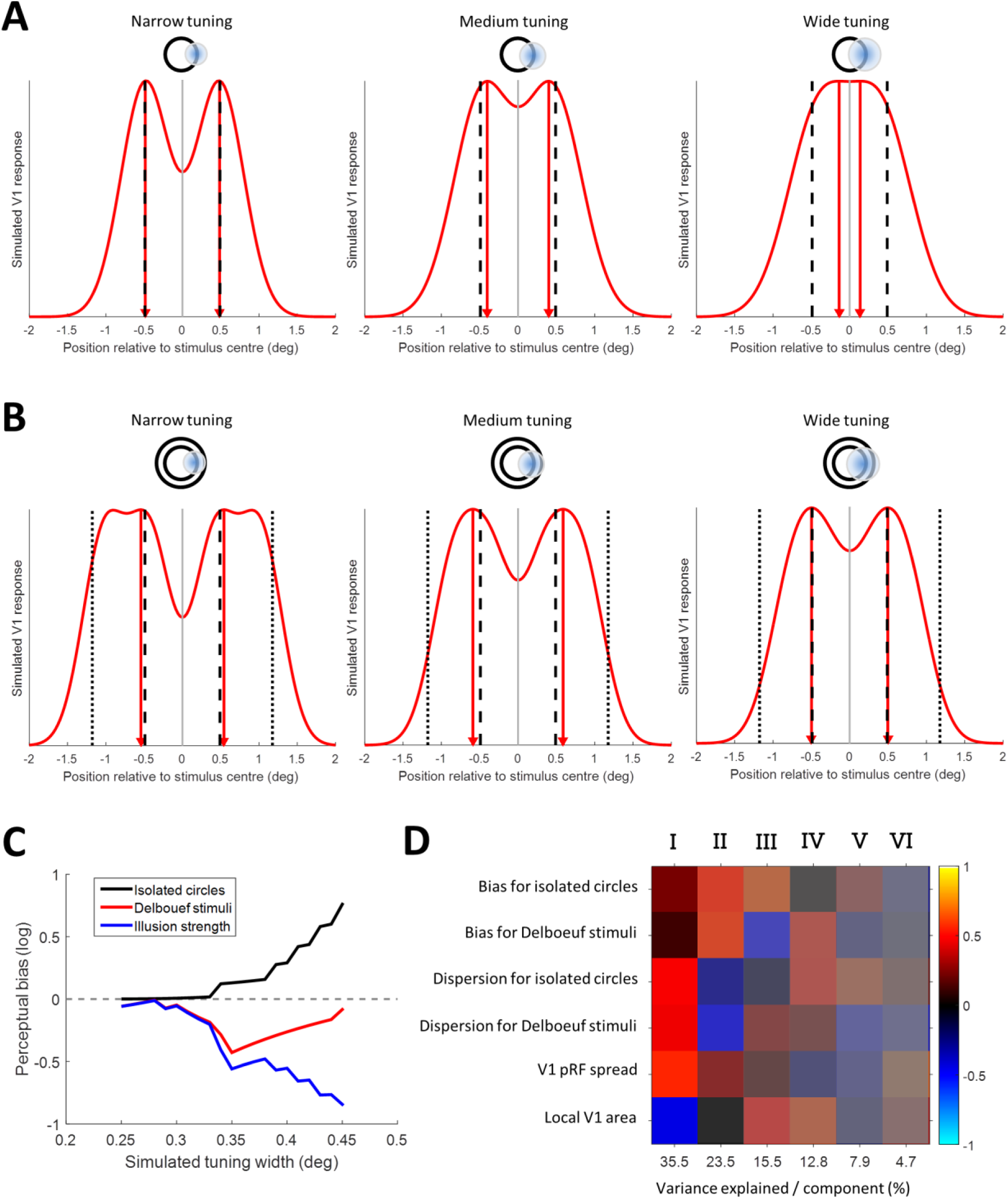
Basic read-out model. **A-B**. Simulated activity profiles for isolated circle (**A**) and Delboeuf stimuli (**B**) were generated by passing the actual location of the stimulus edges through a bank of Gaussian filters covering the stimulus locations. The red curves indicate the output of the filter bank as an simulation of stimulus-related population activity in V1. The separation between the two peaks is an estimate of stimulus size (red triangles and dashed red lines). The vertical black lines denote the actual position of the edges of the target stimulus (dashed) and the inducer annulus in the Delboeuf stimuli (dotted). Simulated neuronal tuning width increases from left to right. This is also illustrated by the schematic diagram above each graph representing the stimulus and an example receptive field. **C**. The simulated perceptual biases for the two stimulus types and the relative illusion strength plotting for a range of tuning widths. Black: isolated circles. Red: Delboeuf stimuli. Blue: Relative illusion strength. See Figure 1D for comparison. **D**. Principal component analysis on a data set combining the biases for isolated circles and Delboeuf stimuli, their respective dispersions (i.e. discrimination acuity), V1 pRF spread, and local V1 surface area (relative to the area of the whole cortical hemisphere). Columns indicate the six principal components. The numbers on the x-axis show the percentage of variance explained by each component.

To quantify the perceptual biases predicted by this model, we simulated perceptual judgments for both stimulus types and across a range of neuronal tuning widths. The relationship between increasing tuning width and perceptual biases parallels that between empirically observed perceptual biases and stimulus eccentricity (Figure 3C). Size estimates at very small receptive fields (and thus lower eccentricity) are largely accurate but apparent size becomes increasingly smaller than the physical stimulus as tuning width increases. Estimates for the Delboeuf stimuli are generally larger than the physical target. However, as tuning width increases estimates again become smaller. Thus, a large part of the difference in perceptual quality between these two stimulus types may be simply due to the physical difference between them, and the corresponding representation within a population of receptive fields, rather than a more complex interaction between the target and the surrounding annulus. Finally, our *relative* illusion strength is the difference between biases for the two stimulus types. As tuning width increases, this measure in turn becomes larger, just as it does for the empirical data in Figure 2D.

One caveat to this model is that the magnitude of simulated biases is a lot larger than those we observed empirically. This may indicate additional processes involved in calibrating the size judgment. However, it may also be simplistic to infer size from the actual activity peaks. The actual read-out process may instead calculate a confidence range that accounts for the whole function describing the activity profile^37^. The exact relationship between pRF spread and neuronal receptive field size is also unknown. Estimates of pRF spread from fMRI data must aggregate the actual *sizes* of neuronal receptive fields, but also the *range* of center positions of all the receptive fields in the voxel, and their *local* positional scatter within this range. In addition, extra-classical receptive field interactions, response nonlinearities^38^, and non-neuronal factors like hemodynamic effects, fixation stability, and head motion must also contribute to some extent. While the simulated tuning widths in our model probably roughly correspond to *neuronal* receptive field size in V1 within our eccentricity range (see e.g. Figure 9B in ref.^31^), an aggregate of the different factors contributing to pRF spread may thus be more appropriate. However, at least qualitatively the relationship between perceptual biases and tuning width parallels the empirical pattern of perceptual biases and pRF spread estimates that we found.

Each row is one of the six variables. The color of each cell denotes the sign (red: positive, blue: negative) and magnitude of how much each variable contributes to each principal component. The saturation denotes the amount of variance explained by each principal component.

### Dissociation between basic perceptual bias and contextual illusions

At the smallest eccentricity of 1.96°, raw perceptual biases for isolated circles were correlated with local V1 area but this pattern was not evident when data were pooled across eccentricity (r=-0.0, p=0.9649; Figure 2F and Supplementary Figure S4A). With Delboeuf stimuli, raw perceptual biases (relative to the central reference) did not correlate with local V1 area at any eccentricity (r=-0.09, p=0.3152; Figure 2G and Supplementary Figure S4B). Because previous research has compared perception to the macroscopic surface area of the entire central portion of V1^8,9,11–15,17^, we further calculated the overall surface area of V1, representing each visual field quadrant between an eccentricity of 1° and 9°. This showed a similar relationship with perceptual biases as local V1 area at the innermost eccentricity (Supplementary Figure S5; isolated circles: r=0.27, p=0.0029; Delboeuf stimuli: r=-0.08, p=0.3968). These results suggest that the variability in perceptual biases is largely driven by differences in cortical magnification for the central visual field: For our innermost eccentricity the relationship between surface area and perceptual measures was always strongest. This variability in central V1 area may thus dominate measurements of the whole quadrant. However, the macroscopic surface area should also be a more stable measure than the area of small local cortical patches. The local surface area and overall area of quadrant maps were very strongly correlated (area relative to whole cortex: r=0.54, p<0.0001; absolute surface area: r=0.54, p<0.0001; see Supplementary Figure S6 for plots separated by eccentricity). Therefore, the macroscopic V1 surface area is a close proxy for local variations in cortical magnification.

In an indirect replication of our earlier findings^8,9,11^, we also observed an inverse relationship between the *relative* strength of the Delboeuf illusion (the difference in perceptual bias measured for the two stimulus types) and V1 surface area. Again this was only significant at the smallest eccentricity and not when data were pooled across eccentricities (r=-0.06, p=0.5066; Figure 2H and Supplementary Figure S5C) but it was significant for the overall area of the quadrant map (r=-0.28, p=0.0017). The relative illusion strength (and thus presumably the Delboeuf illusion itself) is the *difference* in apparent size between these stimuli at the *same location*. This measure could be partially independent of pRF spread as it may instead be related to long-range horizontal connections that exceed the voxel size and that mediate the contextual interaction between target and surround.

Under the hypothesis that surface area predicts illusion strength, the bias induced by the illusion differs mechanistically from basic perceptual biases. Both isolated circles (Figure 2C) and Delboeuf stimuli (Figure 2D) were perceived as smaller when pRFs were larger. However, *at the same location* Delboeuf stimuli were nonetheless seen as larger than isolated circles. Even though the apparent size of both isolated circles and Delboeuf stimuli was linked to pRF spread – consistent with the basic read-out model – the *difference* between these biases was also modestly correlated with the area of central V1. The illusion effect may be modulated by cortical distance, possibly via lateral intra-cortical connections^1,10^, rather than pRF spread. We conjectured previously that the illusion could arise due to long-range connectivity between V1 neurons encoding the target circle and the surrounding context. Thus the illusion may be weaker when V1 surface area (and thus cortical distance) is larger^8–11^. In contrast, basic perceptual biases for any stimulus seem to be linked to the coarseness (pRF spread) of the retinotopic stimulus representation itself, which relates to neuronal receptive field sizes and their local positional scatter.

This interpretation may seem to contradict previous findings that pRF spread is inversely related to V1 surface area^17,35^. However, there is considerable additional unexplained variance to this relationship. To further disentangle the potential underlying factors, we conducted a principal component analysis on a multivariate data set, including z-standardized raw biases for isolated circles and Delboeuf stimuli, the respective dispersions of these distributions (as an indicator of discrimination thresholds), and pRF spread estimates as well as local surface area at corresponding locations in V1. The first four components explained over 87% of the variance (Figure 3D). The first component suggests a positive relationship between pRF spread and dispersion and a negative relationship with V1 area. This supports earlier findings linking pRF spread and cortical magnification to acuity^16,17^. There is however little relation between these measures and perceptual biases. The second component shows a positive relationship between biases for both stimulus types and pRF spread, which reflects our present results (Figure 2C–D). In contrast, the third component involves a negative correlation between biases for the two stimulus types and a positive link between raw biases for isolated circles and V1 area. This may explain the negative correlation between relative illusion strength and V1 area (Supplementary Figure S4C). The fourth component involves a positive link between dispersion for isolated circles and biases for Delboeuf stimuli and also with V1 area. This resembles our earlier findings for orientation processing that also suggest a link between discrimination thresholds for isolated grating stimuli, the strength of the contextual tilt illusion, and V1 surface area^11^.

Taken together, these results indicate that different mechanisms influence apparent size: both isolated circles and Delboeuf stimuli generally appear smaller (relative to the central reference) when pRFs are large, as predicted by the read-out model. However, variability in cortical surface area (and thus the scale required of intra-cortical connections) also seems to be an important factor in the illusory modulation of apparent size. Because our task estimates perceptual biases under either condition relative to a *constant* reference, it was uniquely suited to reveal dissociations between these effects. A more traditional task in which reference stimuli are presented at matched locations/eccentricities would be insensitive to this difference.

### Heterogeneity in perceptual biases has central origin

Naturally, the spatial heterogeneity in perceptual biases could possibly arise from factors prior to visual cortical processing, like small corneal aberrations, inhomogeneity in retinal organization, or the morphology of retinotopic maps in the lateral geniculate nucleus. We tested this possibility in a behavioral control experiment in which we measured perceptual biases while we presented the stimuli either binocularly or dichoptically to the left and right eye. There was a close correspondence between biases measured with either eye (r=0.51, p=0.0103; Figure 4). Thus at least a large part of the variance in perceptual biases must arise at a higher stage of visual processing where the input from both eyes has converged, such as the binocular cells in V1.

**Figure 4.**
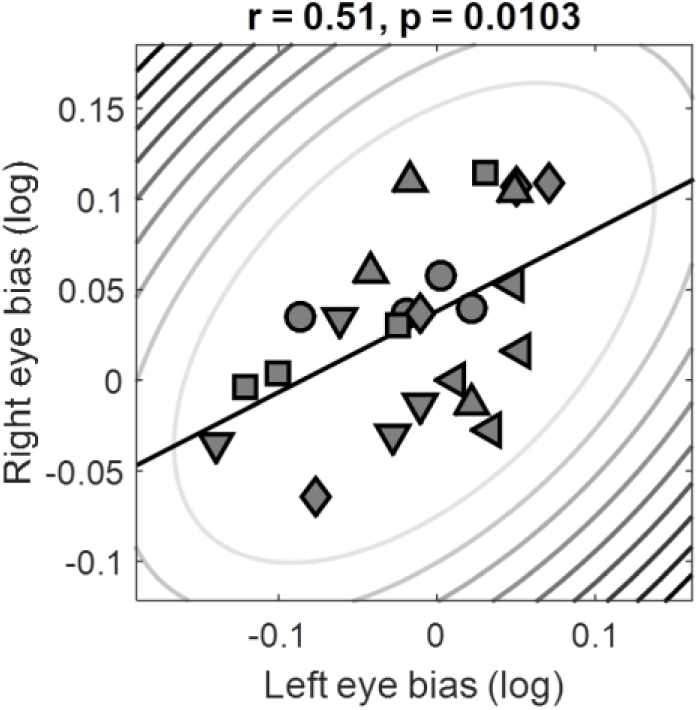
Perceptual biases measured under dichoptic presentation. Biases measured with stimuli in the left eye plotted against those measured in the right eye. Symbols denote individual observers. Elliptic contours denote the Mahalanobis distance from the bivariate mean. The straight, black lines denote the linear regression.

## Discussion

Our experiments show that when the spatial tuning of neuronal populations in V1 is coarse, visual objects are experienced as *smaller*. These findings support the hypothesis that object size is inferred by decision-making processes from the retinotopic representations in V1^1^ and possibly other early visual regions. Our results are therefore consistent with previous reports of a neural signature of apparent size in V1 responses^2–7^. Here we demonstrate that raw perceptual biases are correlated with spatial tuning of neuronal populations in V1. This provides strong evidence that the representation in early visual cortex is indeed used for perceptual decisions about stimulus size, because the biases we observed were independent from contextual or top-down modulation of early visual cortex. Considering that perceptual biases correlate with cortical measures acquired a year later and under completely independent conditions (MRI scanner vs behavioral testing room) we posit that this link between cortical measures and perception is a stable feature of the human visual system.

We have formulated a basic read-out model that samples the activity in early retinotopic maps to infer stimulus size. This model predicts the relationship we observed between eccentricity, pRF spread, and raw perceptual biases measured behaviorally. This was true both for simple, isolated circle stimuli and the Delboeuf stimuli in which the target was surrounded by an annulus. Taking advantage of our unique task design, we further demonstrate that processes related to basic perceptual biases are dissociable from contextual effects, like the Delboeuf illusion: While raw perceptual biases of object size are explained by pRF spread, the local surface area (a measure of cortical distance) of at least the part of V1 representing the very central visual field also explains some variance in contextual modulation of apparent size in these illusions. This underlines the need for a greater understanding of how cortical distance relates to pRF spread. Note however that we only calculated the *relative* strength of the Delboeuf illusion based on the biases measured for the two stimulus types, isolated circles and the contextual stimulus including an annulus. It is possible that this prediction does not fully account for the illusion strength one would measure in more standard procedures.

An alternative explanation to our read-out model is that higher-level decoding mechanisms misestimate the size of the stimulus because they are inadequately calibrated to idiosyncratic differences in cortical magnification^20^. This would cause a residual error between the grossly calibrated read-out that may be reflected in perceptual judgments. This explanation however does not explain why perceptual biases are consistently *reductions* in apparent size. In contrast, our basic read-out model fully accounts for this pattern of results. However, we do not wish to rule out the calibration error hypothesis entirely and in fact a hybrid of the two is certainly possible. In particular, in foveal vision, where individual differences in cortical magnification are far more pronounced^32^, calibration errors are likely. Moreover, the perceptual biases predicted by our basic model are considerably larger than those we observed empirically (even though the relationship with eccentricity parallels the observed data). This finding is consistent with a calibration mechanism that compensates for the incorrect estimation based on basic read-out of the activity in V1.

Naturally, absence of evidence is not evidence of absence. It is entirely possible that some modulations of apparent size are mediated solely by higher-level brain regions but are not represented in early visual cortex. These higher-level areas are of course likely to be involved even in our experiments. Nonetheless, differences in stimulus representations caused by idiosyncratic spatial tuning should be inherited by areas downstream the visual hierarchy, such as V2 and V3. In fact, we observed similar correlations between perceptual biases and pRF spread in V2 and V3. This is unsurprising given the pRF spreads across these early visual areas are also strongly correlated (though interestingly the surface areas of these regions are far less linked^34^). Therefore signals in these regions may also be used for perceptual judgments. However, V1 would be a natural candidate for size estimates given it is the region with the smallest receptive fields and thus the finest spatial resolution. Future research must explore the neural substrate of size judgments, in particular with regard to where in the brain the sensory input is integrated into a perceptual decision^1^. Interestingly, topographically organized tuning for visual object size has recently been reported in parietal cortex^39^. Brain stimulation techniques may help to understand the causal link between early visual cortex and higher decision-making centers.

Our present findings imply that measurements of functional architecture in early sensory cortex can predict individual differences not only of objective discrimination abilities but also our subjective experience of the world. Theoretically, the principle discovered here should also apply to other sensory modalities, such as tuning for auditory frequency or tactile position, and may generalize to more complex forms of tuning, such as object identity or numerosity^40^.

## Acknowledgements

This research was supported by an ERC Starting Grant (310829) to DSS, a Research Fellowship by the Deutsche Forschungsgemeinschaft (Ha 7574/1-1) to BdH, a UCL Graduate School Bridging Fund and a Wellcome Trust Institutional Strategic Support Fund to CM, and a UK MRC Career Development Award (MR/K024817/1) to JAG.

## Methods

### Observers

The authors and several naïve observers participated in these experiments. All observers were healthy and had normal or corrected-to-normal visual acuity. All observers gave written informed consent and procedures were approved by the UCL Research Ethics Committee.

Ten observers (4 authors; 3 female; 2 left-handed; ages 24-37) took part in both the first behavioral experiment measuring perceptual biases at 3 eccentricities and in the fMRI retinotopic mapping experiment (henceforth, *size eccentricity bias* experiment). An additional 3 observers (1 female; all right-handed; ages 20-25) took part in behavioral experiments only but could not be recruited for the fMRI sessions (which commenced several months later and were conducted over the course of a year). These fMRI data form also part of a different study investigating the inter-session reliability of pRF analysis that we are preparing for a separate publication. Nine of the observers from the size eccentricity bias experiment (3 authors; 3 female; 1 left-handed; ages 25-37 at second test) took part in an additional behavioral experiment approximately one year after the first measuring perceptual biases in size perception (*long-term bias reliability*). Four observers (4 authors, 1 female, all right-handed; ages 33-38) participated in another experiment two years after the main experiment to again assess the reliability of bias estimates and also test a greater range of eccentricities (*size far eccentricity bias*). Six observers (5 authors; 2 female; all right-handed; ages 21-36) participated in the dichoptic control experiment (*dichoptic bias*).

### General psychophysical procedure

Observers were seated in a dark, noise-shielded room in front of a computer screen (Samsung 2233RZ) using its native resolution of 1680 x 1050 pixels and a refresh rate of 120Hz. Minimum and maximum luminance values were 0.25 and 230cd/m^2^. Head position was held at 48cm from the screen with a chinrest. Observers used both hands to indicate responses by pressing buttons on a keyboard.

The dichoptic control experiment took place in a different testing room, using an Asus VG278 27” LCD monitor running its native resolution of 1920 x 1080 pixels and a refresh rate of 120Hz. Minimum and maximum luminance values were 0.16 and 100cd/m^2^, with a viewing distance of 60 cm ensured with a chinrest. To produce dichoptic stimulation observers wore nVidia 3D Vision 2 shutter goggles synchronized with the refresh rate of the monitor. Frames for left and right eye stimulation thus alternated at 120Hz.

### Multiple Alternatives Perceptual Search (MAPS) procedure

To estimate perceptual biases efficiently at four visual field locations we developed the MAPS procedure. This is a matching paradigm using analyses related to reverse correlation or classification image approaches^22,23^ that seeks to directly estimate the points of subjective equality, whilst also allowing an inference of discrimination ability.

#### Stimuli

All stimuli were generated and displayed using MATLAB (The MathWorks Inc., Natick, MA) and the Psychophysics Toolbox version 3^41^. The stimuli in all the size discrimination experiments comprised light grey (54cd/m^2^) circle outlines presented on a black background. Each stimulus array consisted of five circles (Figure 1B). One, the *reference*, was presented in the center of the screen and was always constant in size (diameter: 0.98° visual angle). The remaining four, the *targets*, varied in size from trial to trial and independently from each other. They were presented at the four diagonal polar angles and at a distance of 3.92° visual angle from the reference, except for the size eccentricity bias experiment where target eccentricity could be 1.96°, 3.92°, or 7.84° visual angle and the size far eccentricity bias experiment where there were two additional eccentricities in the periphery (11.76° and 15.68°). To measure the bias under the Delboeuf illusion, a larger inducer circle (diameter: 2.35°) surrounded each of the four target circles (but not the reference) to produce a contextual modulation of apparent size.

In all experiments, the independent variable (the stimulus dimension used to manipulate each of the targets) was the binary logarithm of the ratio of diameters for the target relative to the reference circles. In the size eccentricity bias experiment only, the sizes of the four targets were drawn without replacement from a set of fixed sizes (0, ±0.05, ±0.1, ±0.15, ±0.2, ±0.25, ±0.5, ±0.75, or ±1 log units). Thus, frequently there was no “correct” target to choose from. Because this made the task feel quite difficult for many observers, in subsequent experiments (long-term reliability and dichoptic bias) we decided to select a random subset of three targets from a Gaussian noise distribution centered on 0 (the size/orientation of the reference) while one target was correct, i.e. it was set to 0. The standard deviation of the Gaussian noise was 0.3 log units for size discrimination experiments.

#### Tasks

Each trial started with 500ms during which only a fixation dot (diameter: 0.2°) was visible in the middle of the screen. This was followed by presentation of the stimulus array for 200ms after which the screen returned to the fixation-only screen. Observers were instructed to make their response by pressing the F, V, K, or M button on the keyboard corresponding to which of the four targets appeared most similar to the reference. After their response a “ripple” effect over the target they had chosen provided feedback about their response. In the size discrimination experiments this constituted three 50ms frames in which a circle increased in diameter from 0.49° in steps of 0.33° and in luminance.

Moreover, the color of the fixation dot also changed during these 150ms to provide feedback about whether the behavioral response was correct. In the size eccentricity bias experiment, the color was green and slightly larger (0.33°) for correct trials and red for incorrect trials. In all later experiments, we only provided feedback on *correct* trials. This helped to reduce the anxiety associated with large numbers of incorrect trials that are common in this task: Accuracy was typically around 45-50% correct. Even though this is well in excess of chance performance of 25% it means that observers would frequently make mistakes. See Supplementary Information for further details on the task procedure.

Experimental runs were broken up into blocks of 20 trials. After each block there was a resting break. A message on the screen reminded observers of the task and indicated how many blocks they had already completed. Observers initiated blocks with a button press.

*Size eccentricity bias experiment:* Observers were recruited for two sessions on separate days. In each session they performed six experimental runs, three with only circles and three with the Delboeuf stimuli. Each run tested one of the three target eccentricities. Trials with different eccentricities were run in separate blocks to avoid confounding these measurements with differences in attentional deployment across different eccentricities. There were 10 blocks per experimental run. In the *size far eccentricity bias* experiment we only tested observers in one session on the five target eccentricities.

*Long-term bias reliability experiment:* Half of the experimental runs observers performed measured their baseline biases. The other half of the runs contained artificially induced biases: two of the four targets were altered subtly: one by adding and one by subtracting 0.1 log units. Which two targets were altered was counterbalanced across observers, as was the order of experimental runs. Observers were recruited for only one session comprising four runs (two with artificial bias) plus an additional run measuring biases for the Delboeuf stimuli. There were 10 blocks per experimental run. Only the results of the baseline biases (i.e. without artificially induced bias) are presented in the present study. The remainder of these experiments form part of another study and will be presented elsewhere.

*Dichoptic bias experiment:* There were three experimental conditions in this experiment. By means of shutter goggles the stimulus arrays could be presented dichoptically, either binocularly or monocularly to either eye. To aid stereoscopic fusion we additionally added 5 concentric squares surrounding the stimulus arrays (side length: 8.1-10.5° in equal steps). The three experimental conditions were randomly interleaved within each experimental run. There were 34 blocks per run; however, in this experiment each block comprised only 12 trials. Observers performed two such runs within a single session.

#### Analysis

To estimate perceptual biases we fit a model to predict a given observer’s behavioral response in each trial (Figure 1C). For each target stimulus location a Gaussian tuning curve denoted the output of a “neural detector”. The detector producing the strongest output determined the predicted choice. The model fitted the peak location (μ) and dispersion (σ) parameters of the Gaussian tuning curves that minimized the prediction error across all trials. Model fitting employed the Nelder-Mead simplex search optimization procedure^42^. We initialized the μ parameter as the mean stimulus value (offset in logarithmic size ratio from 0) whenever a given target location was chosen *incorrectly*. We initialized the σ parameter as the standard deviation across all stimulus values when a given target location was chosen. The final model fitting procedure however always used *all* trials, correct and incorrect.

### Retinotopic mapping experiment

The same ten observers from the size eccentricity bias experiment participated in two sessions of retinotopic mapping in a Siemens Avanto 1.5T MRI scanner using a 32-channel head coil located at the Birkbeck-UCL Centre for Neuroimaging. The front half of the coil was removed to allow unrestricted field of view leaving 20 channels. Observers lay supine and watched the mapping stimuli, which were projected onto a screen (resolution: 1920 x 1080) at the back of the bore, via a mirror mounted on the head coil. The viewing distance was 68cm.

We used a T2*-weighted multiband 2D echo-planar sequence^43^ to acquire 235 functional volumes per pRF mapping run and 310 volumes for a run to estimate the hemodynamic response function (HRF). In addition, we collected a T1-weighted anatomical magnetization-prepared rapid acquisition with gradient echo (MPRAGE) scan with 1 mm isotropic voxels (TR=2730ms, TE=3.57ms) using the full 32-channel head coil.

The method we used for analyzing pRF^31^ data has been described previously^32,33^. We used a combined wedge and ring stimulus that contained natural images that changed twice a second (see Supplementary Information for further details on the design of the mapping experiments). The MATLAB toolbox (available at http://dx.doi.org/10.6084/m9.figshare.1344765) models the pRF of each voxel as a two-dimensional Gaussian in visual space and incorporates the hemodynamic response function measured for each individual observer. It determines the visual field location (x and y in Cartesian coordinates) and the spread (standard deviation) of the pRF plus an overall response amplitude.

#### Stimuli and task

A polar wedge subtending a polar angle of 12° rotated in 60 discrete steps (one per second) around the fixation dot (diameter: 0.13° surrounded by a 0.25° annulus where contrast ramped up linearly). A ring expanded or contracted, both in width and overall diameter, in 36 logarithmic steps. The maximal eccentricity of the wedge and ring was 8.5°. There were 3 cycles of wedge rotation and 5 cycles of ring expansion/contraction. Each mapping run concluded with 45s of a fixation-only period. At all times a low contrast ‘radar screen’ pattern (Figure 2A) was superimposed on the screen to aid fixation compliance.

The wedge and ring parts contained colorful natural images (Figure 2A) from Google Image search, which changed every 500ms. They depicted outdoor scenes (tropical beaches, forests, mountains, and rural landscapes), faces, various animals, and pictures of written script (228 images in total). One picture depicted the ‘Modern Anderson’ clan tartan. These pictures were always rotated in accordance with the current orientation of the wedge. Observers were asked to fixate a fixation dot at all times. With a probability of 0.03 every 200ms the black fixation dot would change color for 200ms to one of the primary and complementary colors or white followed by another 200ms of black. Observers were asked to tap their finger when the dot turned red. To also maintain attention on the mapping stimulus they were asked to tap their finger whenever they saw the tartan image.

In alternating runs the wedge rotated in clockwise and counterclockwise directions, while the ring expanded and contracted, respectively. In each session we collected six such mapping runs and an additional run to estimate the hemodynamic response function. The latter contained 10 trials each of which started with a 2s sequence of four natural images from the same set used for mapping. These were presented in a circular aperture centered on fixation with radius 8.5° visual angle. This was followed by 28s of the blank screen (fixation and radar screen only).

#### Preprocessing and pRF modeling

Functional MRI data were first preprocessed using SPM8 (Wellcome Trust Centre for Neuroimaging, London, http://www.fil.ion.ucl.ac.uk/spm/software/spm8). The first 10 volumes were removed to allow the signal to reach equilibrium. We performed intensity bias correction, realignment and unwarping, and coregistration of the functional data to the structural scan, all using default parameters. We used FreeSurfer (https://surfer.nmr.mgh.harvard.edu/fswiki) for automatic segmentation and reconstruction to create a three-dimensional inflated model of the cortical surfaces for the grey-white matter boundary and the pial surface, respectively^44,45^. We then projected the functional data to the cortical surface by finding for each vertex in the surface mesh the median position between the grey-white matter and pial surfaces in the functional volume. All further analyses were performed in surface space.

We applied linear detrending to the time series from each vertex in each run and then z-standardized them. Alternating pRF mapping runs (i.e. those sharing the same stimulus directions – clockwise/expanding and counterclockwise/contracting) were averaged. These two average runs were then concatenated. We further divided the HRF run into the 10 epochs and averaged them. Only vertexes for which the average response minus the standard error in the first half of the trial was larger than zero were included. The HRFs for these vertices were than averaged and we fit a two-gamma function with four free parameters: the amplitude, the peak latency, the undershoot latency, and the ratio amplitude between peak and undershoot.

Population receptive field analysis was conducted in a two-stage procedure. First, a coarse fit was performed on data smoothed with a large kernel on the spherical surface (FWHM=5mm). We performed an extensive grid search on every permutation of 15 plausible values for x and y, respectively, and a range of pRF spreads from 0.18° to 17° in 34 logarithmic steps (0.2 in binary logarithm). For each permutation we generated a predicted time series by calculating the overlap between a two-dimensional Gaussian pRF profile and a binary aperture of the mapping stimulus for every volume. This time series was then z-standardized and convolved with the subject-specific HRF. The grid search is a very fast operation that computes the set of three pRF parameters that produce the maximal Pearson correlation between the predicted and observed time series for the whole set of search grid parameters and all vertices. This was followed by the slow fine fit. Here we used the parameters identified by the coarse fit to seed an optimization algorithm^42,46^ on a vertex by vertex basis to refine the parameter estimates by minimizing the squared residuals between the predicted and observed time series. This stage used the unsmoothed functional data and also included a fourth amplitude parameter to estimate response strength. Finally, the estimates parameter maps were also smoothed on the spherical surface with a modest kernel (FWHM=3mm).

#### Analysis of functional cortical architecture

We next delineated the early visual regions (specifically V1) manually based on reversals in the polar angle map and the extent of the activated portion of visual cortex along the anterior-posterior axis. We then extracted the pRF parameter data separately from each visual field quadrant represented in V1. Data were divided into eccentricity bands 1° in width starting from 1° eccentricity up to 9°. For each eccentricity band we then calculated mean pRF spread and the sum of surface area estimates. For pRF spread we used the raw, unsmoothed pRF spread estimates produced by our fine-fitting procedure. However, the quantification of surface area requires a smooth gradient in the eccentricity map without any gaps in the map and with minimal position scatter in pRF positions. Therefore, we used the final smoothed parameter maps for this analysis. The results for pRF spread are very consistent when using smoothed parameter maps but we reasoned that unsmoothed data make fewer assumptions.

To extract the parameters for each stimulus location we fit polynomial functions to the relationship between these binned parameters and eccentricity. For pRF spread we used a first order polynomial (i.e. a linear relationship). For surface area we used a second order polynomial. We then interpolated each person’s pRF spread/surface area at the 12 target locations in the behavioral experiments (that is, 4 stimulus locations and 3 eccentricities. Individual plots for each observer and visual field quadrant are included in the Supplementary Information. We also quantified the macroscopic surface area of each *visual field quadrant* in V1 by summing the surface area between 1° and 9°. This range ensured that artifactual, noisy estimates in the foveal confluence or edge effects well beyond the stimulated region did not introduce spurious differences between individuals. In our main analyses we normalized all surface area measures relative to the whole cortical surface area. However, results are also very consistent for using the square root of the absolute surface area for this analysis.

## Data and materials

Materials and behavioral data: http://dx.doi.org/10.6084/m9.figshare.1579442

Processed pRF data per observer: http://dx.doi.org/10.5281/zenodo.19150

## Supplementary Figures

**Supplementary Figure S1.**
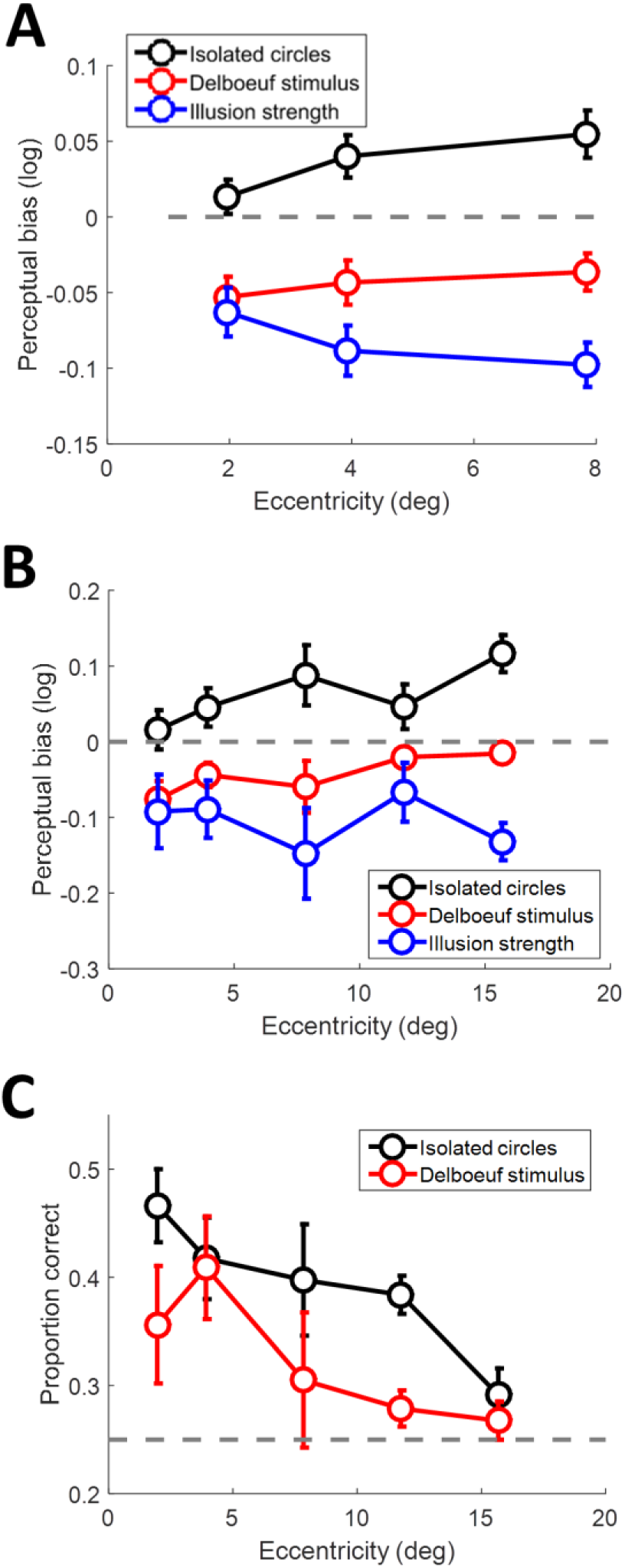
A-B. Average perceptual bias (positive and negative: target appears smaller or larger than reference, respectively) weighted by the acuity (reciprocal of squared dispersion), across individuals plotted against target eccentricity for simple isolated circles (black), contextual Delboeuf stimuli (red), and relative illusion strength (blue), that is, the difference in biases measured for the two stimulus conditions. A. Data from 10 observers in the size eccentricity bias experiment. B. Data from 4 observers in the size far-eccentricity bias experiment. C. Behavioral accuracy on the task in for the 4 observers in the size far-eccentricity bias experiment. Chance was 25% and is noted by the dashed grey line. In all plots, error bars denote ±1 standard error of the mean.

**Supplementary Figure S2.**
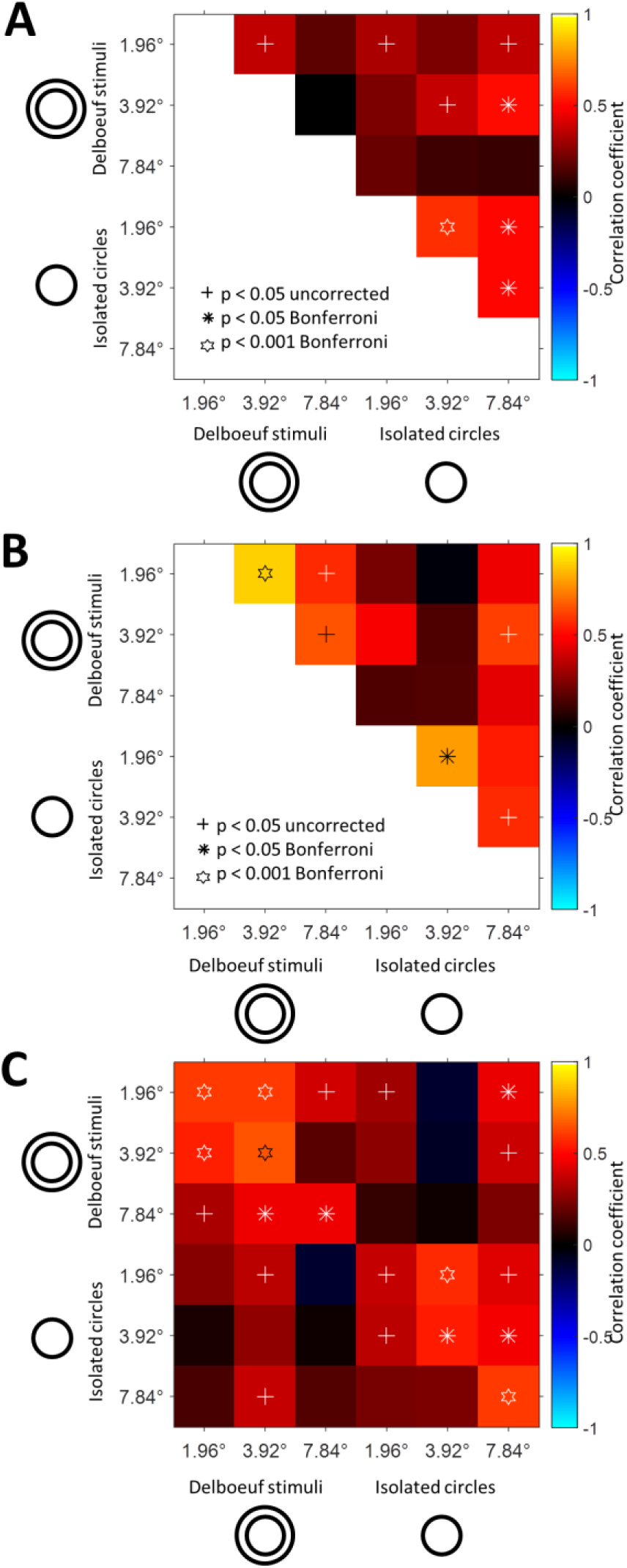
Correlation matrices showing the relationship between the perceptual biases in the two conditions (isolated circles and Delboeuf stimuli) and at the three stimulus eccentricities. **A**. Correlations after removing between-subject variance, i.e. the mean across the biases for the four targets was *subtracted* from each condiion. **B**. Correlations after removing the within-subject variance, i.e. biases were *averaged* across the four targets in each condition. **C**. Correlations between the first and second session of the experiment conducted on different days. All other conventions as in Figure 1E. Note that statistical power in **B** is lower relative to the other figures, because after averaging there is only a quarter of the number of observations.

**Supplementary Figure S3.**
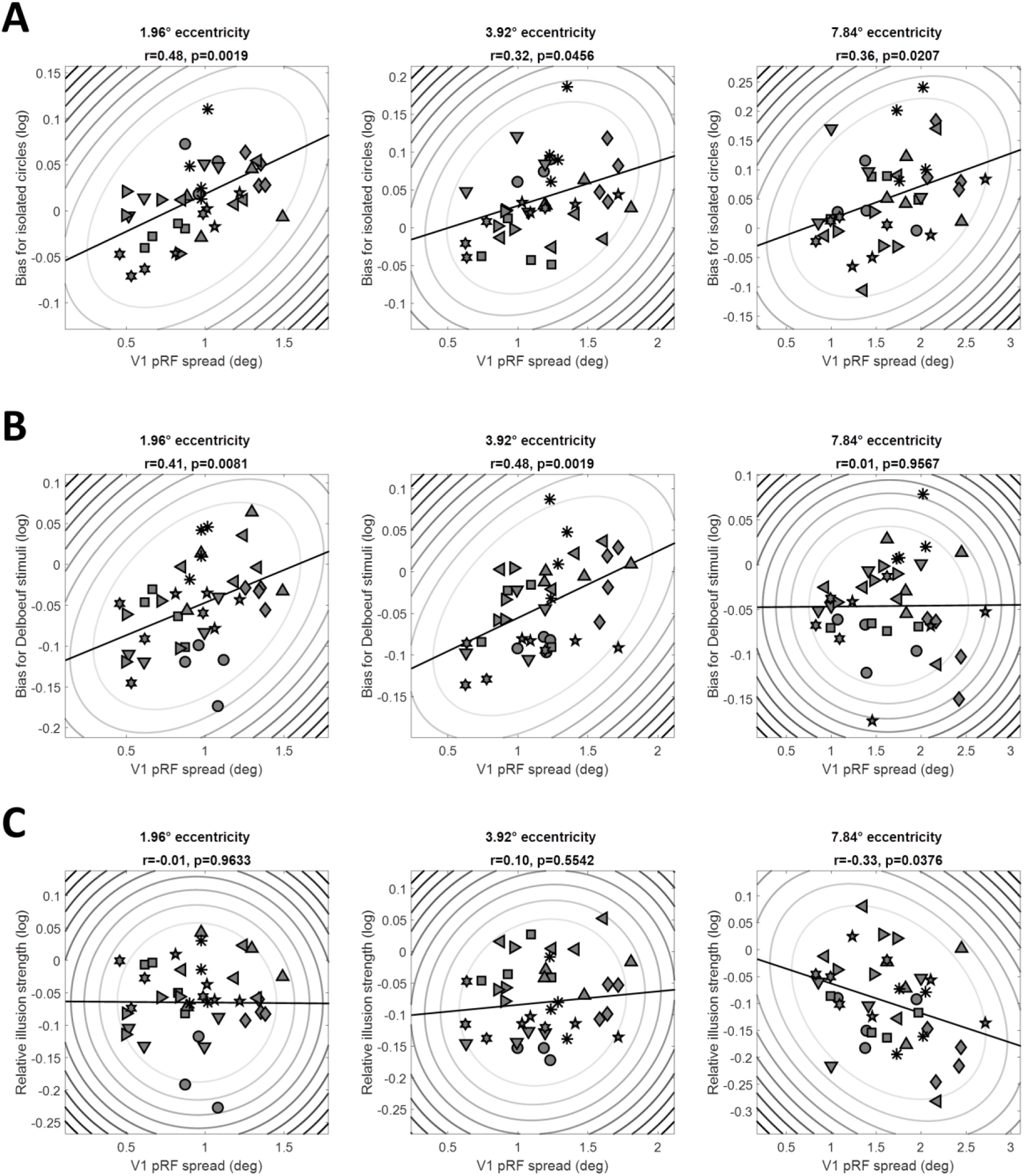
Perceptual biases for isolated circles (**A**), for the Delboeuf stimuli (**B**), and the relative illusion strength (**C**), that is, the bias for Delboeuf stimuli minus the bias for isolated circles, plotted against pRF spread at the corresponding location in V1 for each observer and stimulus location. Columns show data for stimuli at 1.96°, 3.92°, or 7.84° eccentricity. Symbols denote individual observers. Elliptic contours denote the Mahalanobis distance from the bivariate mean. The straight, black lines denote the linear regression.

**Supplementary Figure S4.**
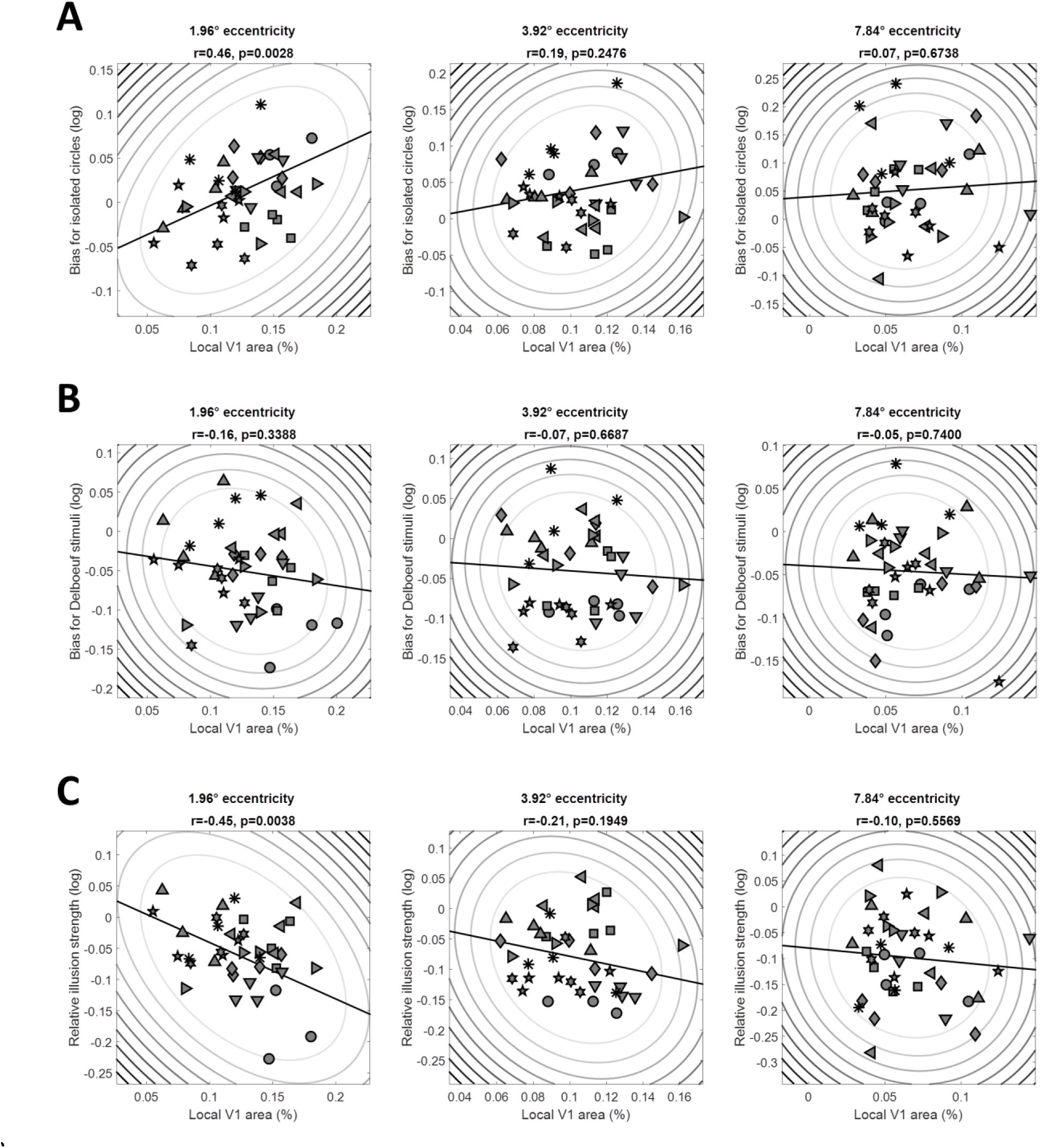
Perceptual biases for isolated circles (**A**), for the Delboeuf stimuli (**B**), and the relative illusion strength (**C**), that is, the bias for Delboeuf stimuli minus the bias for isolated circles, plotted against the surface area of the corresponding location in V1 for each observer and stimulus location (as percentage of the area of the whole cortical hemisphere). Columns show data for stimuli at 1.96°, 3.92°, or 7.84° eccentricity. Symbols denote individual observers. Elliptic contours denote the Mahalanobis distance from the bivariate mean. The straight, black lines denote the linear regression.

**Supplementary Figure S5.**
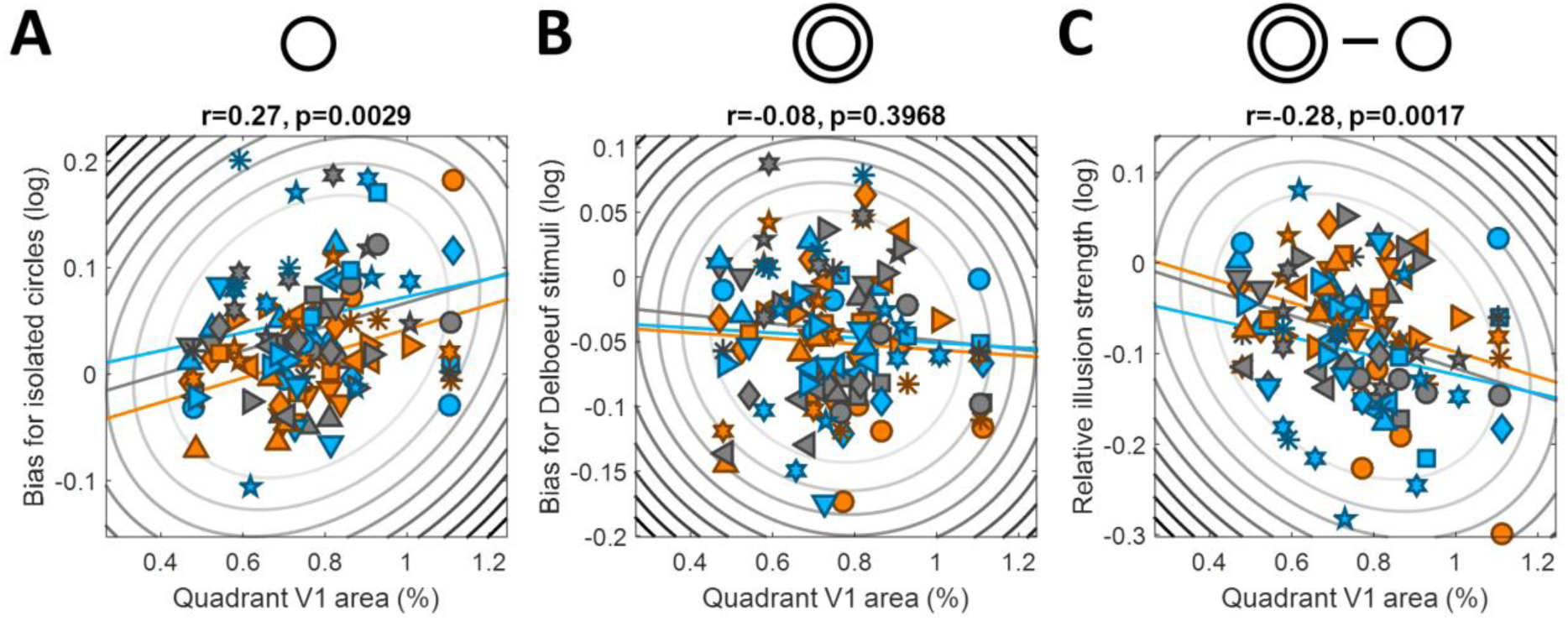
Perceptual biases for isolated circles (**A**), Delboeuf stimuli (**B**), and the relative illusion strength (**C**) plotted against the surface area (as percentage of the whole cortex) for each quadrant map in V1 between 1° and 9° eccentricity and observer. Symbols denote individual observers. Elliptic contours denote the Mahalanobis distance from the bivariate mean. The colored, straight lines denote the linear regression separately for each eccentricity. Colors denote stimuli at 1.96° (orange), 3.92° (grey), or 7.84° (light blue) eccentricity.

**Supplementary Figure S6.**
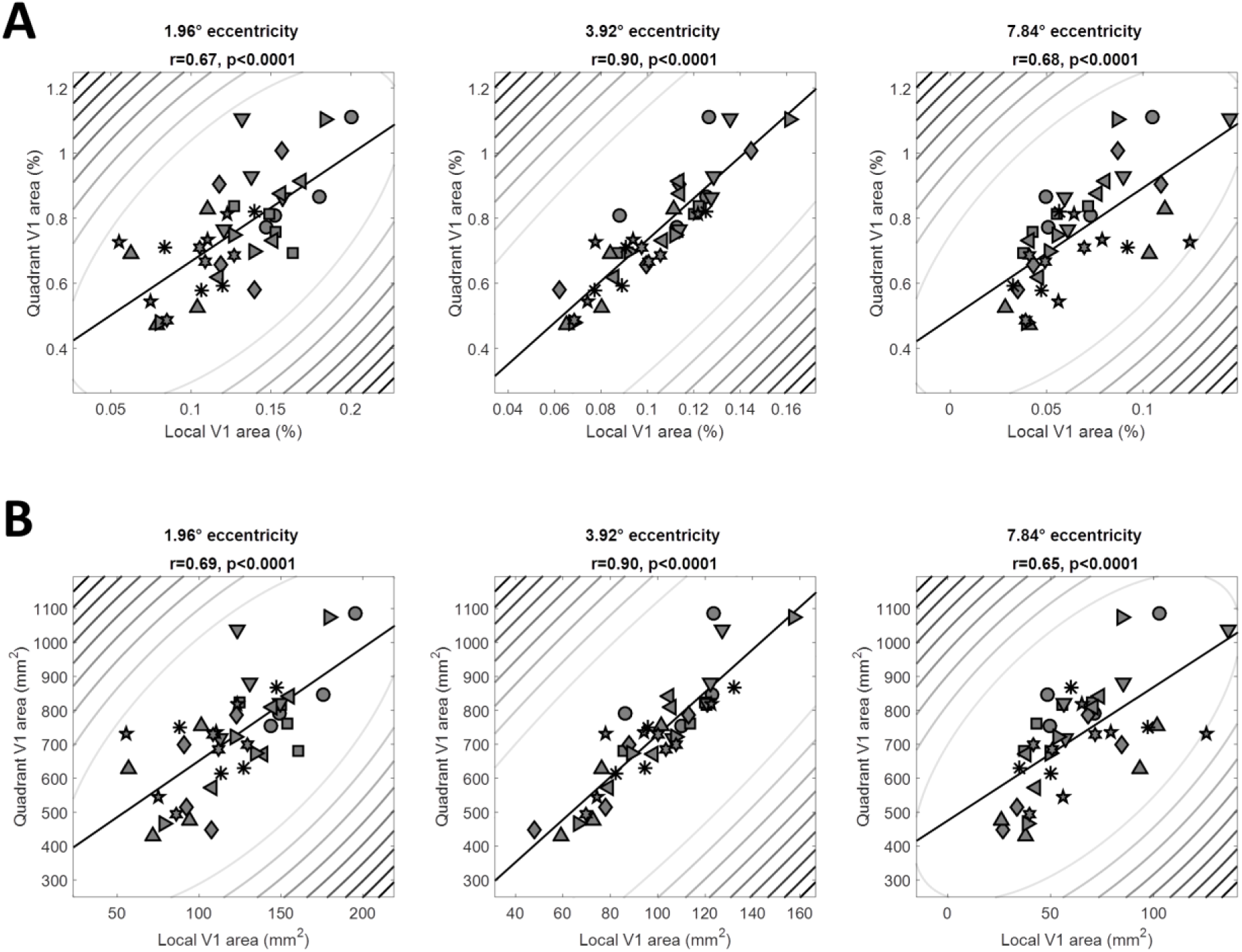
The surface area of the whole quadrant map in V1 between 1° and 9° eccentricity and each observer plotted against the surface area of the corresponding location in V1 for each observer and target stimulus location. Surface area is expressed either as a percentage of the whole cortical hemisphere (**A**) or as absolute area (**B**). Columns show data for stimuli at 1.96°, 3.92°, or 7.84° eccentricity. Symbols denote individual observers. Elliptic contours denote the Mahalanobis distance from the bivariate mean. The straight, black lines denote the linear regression.

## Supplementary Data File caption

Mean V1 pRF spread for the four visual field quadrants (top four panels) and V1 surface area (bottom four panels) plotted for eccentricity bands 1° in width. The vertical dashed red lines indicate the eccentricities of target stimuli in the psychophysical experiments. The solid black lines denote the fitted polynomial functions. Each page shows plots from one observer.

## Supplementary Information

### Reliability of perceptual bias estimates

We further confirmed the reliability of these bias estimates by comparing estimates from two sessions conducted on different days (Supplementary Figure S2C). Moreover, 9 of our observers were tested twice, with approximately one year between sessions. Despite the long time between experiments and variation in the stimulus sampling procedure (see Methods), estimates of perceptual biases at target eccentricity 3.92° (which was common to both experiments) were correlated (r=0.35, p=0.0373, n=36). This correlation was largely driven by the within-subject variance, and was considerably greater after subtracting the mean across the four target locations for every condition (r=0.58, p=0.0002, n=36). In contrast, removing the within-subject variance by averaging bias estimates across the four targets reduced the correlation substantially (r=0.18, p=0.6483, n=36). Finally, 4 observers repeated the experiment two years after the initial experiment, allowing us to compare biases for the three eccentricities tested in the original experiment (n=48). We again found a strong reliability of idiosyncratic biases (r=0.47, p=0.001, n=48; after removing between-subject variance: r=0.71, p<0.001, n=48).

### Intra-individual differences analysis

For each observer we obtained separate measures of perceptual bias and cortical measures corresponding to 12 visual field locations. We then calculated correlations by comparing all locations (120 data points) or across quadrants but separately for each eccentricity (40 data points). Naturally, multiple observations for a given participant are not strictly independent. Therefore, as described in the main text we performed four parallel analyses:

1. *Pooled data (all variance):* The main analyses reported in our study simply show the pooled data without any additional processing. They therefore compare the 120 (or 40, when separating eccentricities) data points with each visual field location as a separate data point (Figure 2 C-E; Figure S3 A-C). This approach is the most inclusive as it incorporates both the within-subject variance (the pattern of variability across visual field locations) as well as the conventional between-subject variance (differences between individual observers that affect all visual field locations in a given observer equally). Our hypothesis that cortical idiosyncrasies in pRF spread/surface area relate to perceptual biases suggests that both between- and within-subject variance should contribute similarly to the correlation.
2. *Within-subject variance only:* We also calculated correlations after removing the between-subject variance by first subtracting the mean of measurements across the four visual field locations from each eccentricity and observer. This way the correlation only takes into account the variability across quadrants within each observer/eccentricity.
3. *Second-level analysis of within-subject variance:* In an alternative analysis using only the within-subject variance we calculated the correlation between the two variables separately for each eccentricity and each observer, and then determined whether the average correlation (after Fisher’s z-transformation to linearize r) is significantly different from zero. However, this approach is comparably underpowered because it relies on only four data points (one per visual field quadrant) for each observer and eccentricity. Thus, each individual correlation coefficient is likely to be skewed by outliers or individual unreliable measurements and this approach is prone to both type I and type II error. 4. *Canonical correlation analysis:* Finally, we used established canonical correlation analysis (*canoncorr* in MATLAB). This allows the use of multiple dimensions (i.e. the different stimulus locations) per data point; however, this procedure overcomplicates the problem because it determines a linear combination of the different dimensions whereas we are strictly interested in comparing the perceptual bias measured at one location to the corresponding cortical measures.

### Power analysis

To confirm the validity of our analysis approach, we conducted simulations to determine its statistical power. In 10,000 simulations we generated random data sets with the same sample sizes and dimensionality of our data to test three situations: A) Complete null hypothesis: the two variables were completely uncorrelated. B) Complete alternative hypothesis: The 120 data points were chosen from the same underlying distribution with a population correlation of 0.3. This is the alternative hypothesis we seek to test in this study, because it assumes that variability in cortical measures (pRF spread or cortical surface area) is directly linked to perceptual biases. C) Between-subject relationship only: two variables of 10 subjects with 12 stimulus locations were correlated (using population correlation of 0.3) but the within-subject variance was random noise (Gaussian noise with 0.5 standard deviations) added to the 4 observations for each observer and eccentricity. This situation assumes the effect is solely driven by the between-subject variance and within-subject variance is merely measurement noise within each observer.

These simulations showed that all four analyses (see *Intra-individual differences analysis*) have nominal levels of false positives (∼5%) when the null hypothesis is true (situation A), except for analysis 2 (within-subject variance only) which somewhat inflates false positive rates to around 9%. Conversely, for situation B when the alternative hypothesis is true and there is a direct relationship between the two variables, analyses 1 and 2 are most sensitive with a statistical power of approximately 92% and 89%, respectively. The other two analyses are far less sensitive (65% power for analysis 3, and 42% power for analysis 4). Finally, in situation C when the relationship is only driven by the between-subject variance, statistical power for analyses 1 is still moderately high (65%) but as expected power for all the analyses is much lower (9% and 10% for analyses 2 and 4, respectively, and at the alpha level of 5% for analysis 3). Thus if our hypothesis of a direct link of the within-subject variability in perceptual biases and V1 measures were untrue and the relationship was mainly driven by between-subject variance, we would have been unlikely to detect any correlations in these control analyses. This is clearly not the case as the pattern of results is qualitatively very similar between the four analyses in most cases – especially the main result comparing pRF spread to perceptual biases of isolate circles is highly significant in all four analyses (Table 1).

## References

1. Schwarzkopf, D. S. Where Is Size in the Brain of the Beholder? Multisensory Res. 28, 285–296 (2015).

2. Murray, S. O., Boyaci, H. & Kersten, D. The representation of perceived angular size in human primary visual cortex. Nat. Neurosci. 9, 429–434 (2006).

3. Fang, F., Boyaci, H., Kersten, D. & Murray, S. O. Attention-dependent representation of a size illusion in human V1. Curr. Biol. 18, 1707–1712 (2008).

4. Sperandio, I., Chouinard, P. A. & Goodale, M. A. Retinotopic activity in V1 reflects the perceived and not the retinal size of an afterimage. Nat. Neurosci. 15, 540–542 (2012).

5. Pooresmaeili, A., Arrighi, R., Biagi, L. & Morrone, M. C. Blood Oxygen Level-Dependent Activation of the Primary Visual Cortex Predicts Size Adaptation Illusion. J. Neurosci. 33, 15999–16008 (2013).

6. Ni, A. M., Murray, S. O. & Horwitz, G. D. Object-Centered Shifts of Receptive Field Positions in Monkey Primary Visual Cortex. Curr. Biol. CB (2014). doi:10.1016/j.cub.2014.06.003

7. He, D., Mo, C., Wang, Y. & Fang, F. Position shifts of fMRI-based population receptive fields in human visual cortex induced by Ponzo illusion. Exp. Brain Res. (2015). doi:10.1007/s00221-015-4425-3

8. Schwarzkopf, D. S., Song, C. & Rees, G. The surface area of human V1 predicts the subjective experience of object size. Nat. Neurosci. 14, 28–30 (2011).

9. Schwarzkopf, D. S. & Rees, G. Subjective size perception depends on central visual cortical magnification in human v1. PloS One 8, e60550 (2013).

10. Song, C. et al. Effective connectivity within human primary visual cortex predicts interindividual diversity in illusory perception. J. Neurosci. Off. J. Soc. Neurosci. 33, 18781–18791 (2013).

11. Song, C., Schwarzkopf, D. S. & Rees, G. Variability in visual cortex size reflects tradeoff between local orientation sensitivity and global orientation modulation. Nat. Commun. 4, 2201 (2013).

12. Genç, E., Bergmann, J., Singer, W. & Kohler, A. Surface Area of Early Visual Cortex Predicts Individual Speed of Traveling Waves During Binocular Rivalry. Cereb. Cortex N. Y. N 1991 (2014). doi:10.1093/cercor/bht342

13. Verghese, A., Kolbe, S. C., Anderson, A. J., Egan, G. F. & Vidyasagar, T. R. Functional size of human visual area V1: A neural correlate of top-down attention. NeuroImage (2014). doi:10.1016/j.neuroimage.2014.02.023

14. Bergmann, J., Genç, E., Kohler, A., Singer, W. & Pearson, J. Neural Anatomy of Primary Visual Cortex Limits Visual Working Memory. Cereb. Cortex N. Y. N 1991 (2014). doi:10.1093/cercor/bhu168

15. Bergmann, J., Genç, E., Kohler, A., Singer, W. & Pearson, J. Smaller Primary Visual Cortex Is Associated with Stronger, but Less Precise Mental Imagery. Cereb. Cortex bhv186 (2015). doi:10.1093/cercor/bhv186

16. Duncan, R. O. & Boynton, G. M. Cortical magnification within human primary visual cortex correlates with acuity thresholds. Neuron 38, 659–671 (2003).

17. Song, C., Schwarzkopf, D. S., Kanai, R. & Rees, G. Neural Population Tuning Links Visual Cortical Anatomy to Human Visual Perception. Neuron (2015). doi:10.1016/j.neuron.2014.12.041

18. Helmholtz, H. Handbuch der Physiologischen Optik. (1867).

19. Newsome, L. R. Visual angle and apparent size of objects in peripheral vision. Percept. Psychophys. 12, 300–304 (1972).

20. Anstis, S. Picturing peripheral acuity. Perception 27, 817–825 (1998).

21. Afraz, A., Pashkam, M. V. & Cavanagh, P. Spatial heterogeneity in the perception of face and form attributes. Curr. Biol. CB 20, 2112–2116 (2010).

22. Abbey, C. K. & Eckstein, M. P. Classification image analysis: estimation and statistical inference for two-alternative forced-choice experiments. J. Vis. 2, 66–78 (2002).

23. Li, R. W., Levi, D. M. & Klein, S. A. Perceptual learning improves efficiency by re-tuning the decision ‘template’ for position discrimination. Nat. Neurosci. 7, 178–183 (2004).

24. Morgan, M., Dillenburger, B., Raphael, S. & Solomon, J. A. Observers can voluntarily shift their psychometric functions without losing sensitivity. Atten. Percept. Psychophys. 74, 185–193 (2012).

25. Morgan, M. J., Melmoth, D. & Solomon, J. A. Linking hypotheses underlying Class A and Class B methods. Vis. Neurosci. 30, 197–206 (2013).

26. Jogan, M. & Stocker, A. A. A new two-alternative forced choice method for the unbiased characterization of perceptual bias and discriminability. J. Vis. 14, 20 (2014).

27. Bedell, H. E. & Johnson, C. A. The perceived size of targets in the peripheral and central visual fields. Ophthalmic Physiol. Opt. J. Br. Coll. Ophthalmic Opt. Optom. 4, 123–131 (1984).

28. Delboeuf, J. Sur une nouvelle illusion d’optique. Acade ¨ Mie R. Sci. Lett. B.-arts Belg. Bull. 24, 545–558 (1892).

29. Abrams, J., Nizam, A. & Carrasco, M. Isoeccentric locations are not equivalent: the extent of the vertical meridian asymmetry. Vision Res. 52, 70–78 (2012).

30. Rubin, N., Nakayama, K. & Shapley, R. Enhanced perception of illusory contours in the lower versus upper visual hemifields. Science 271, 651–653 (1996).

31. Dumoulin, S. O. & Wandell, B. A. Population receptive field estimates in human visual cortex. NeuroImage 39, 647–660 (2008).

32. Schwarzkopf, D. S., Anderson, E. J., Haas, B. de, White, S. J. & Rees, G. Larger Extrastriate Population Receptive Fields in Autism Spectrum Disorders. J. Neurosci. 34, 2713–2724 (2014).

33. Alvarez, I., De Haas, B. A., Clark, C. A., Rees, G. & Schwarzkopf, D. S. Comparing different stimulus configurations for population receptive field mapping in human fMRI. Front. Hum. Neurosci. 9, 96 (2015).

34. Dougherty, R. F. et al. Visual field representations and locations of visual areas V1/2/3 in human visual cortex. J. Vis. 3, 586–598 (2003).

35. Harvey, B. M. & Dumoulin, S. O. The Relationship between Cortical Magnification Factor and Population Receptive Field Size in Human Visual Cortex: Constancies in Cortical Architecture. J. Neurosci. Off. J. Soc. Neurosci. 31, 13604–13612 (2011).

36. Pouget, A., Dayan, P. & Zemel, R. Information processing with population codes. Nat. Rev. Neurosci. 1, 125–132 (2000).

37. Treue, S., Hol, K. & Rauber, H.-J. Seeing multiple directions of motion—physiology and psychophysics. Nat. Neurosci. 3, 270–276 (2000).

38. Kay, K. N., Winawer, J., Mezer, A. & Wandell, B. A. Compressive spatial summation in human visual cortex. J. Neurophysiol. 110, 481–494 (2013).

39. Harvey, B. M., Fracasso, A., Petridou, N. & Dumoulin, S. O. Topographic representations of object size and relationships with numerosity reveal generalized quantity processing in human parietal cortex. Proc. Natl. Acad. Sci. U. S. A. 112, 13525–13530 (2015).

40. Harvey, B. M., Klein, B. P., Petridou, N. & Dumoulin, S. O. Topographic representation of numerosity in the human parietal cortex. Science 341, 1123–1126 (2013).

41. Brainard, D. H. The Psychophysics Toolbox. Spat. Vis. 10, 433–6 (1997).

42. Lagarias, J., Reeds, J., Wright, M. & Wright, P. Convergence properties of the Nelder— Mead simplex method in low dimensions. SIAM J. Optim. 9, 112–147 (1998).

43. Breuer, F. A. et al. Controlled aliasing in parallel imaging results in higher acceleration (CAIPIRINHA) for multi-slice imaging. Magn. Reson. Med. 53, 684–691 (2005).

44. Dale, A. M., Fischl, B. & Sereno, M. I. Cortical surface-based analysis. I. Segmentation and surface reconstruction. NeuroImage 9, 179–194 (1999).

45. Fischl, B., Sereno, M. I. & Dale, A. M. Cortical surface-based analysis. II: Inflation, flattening, and a surface-based coordinate system. NeuroImage 9, 195–207 (1999).

46. Nelder, J. A. & Mead, R. A Simplex Method for Function Minimization. Comput. J. 7, 308–313 (1965).

